# Crosstalk between diverse synthetic protein degradation tags in *Escherichia coli*

**DOI:** 10.1101/136812

**Authors:** Nicholas C. Butzin, William H. Mather

## Abstract

Recently, a synthetic circuit in *E. coli* demonstrated that two proteins engineered with LAA tags targeted to the native protease ClpXP are susceptible to crosstalk due to competition for degradation between proteins. To understand proteolytic crosstalk beyond the single protease regime, we investigated in *E. coli* a set of synthetic circuits designed to probe the dynamics of existing and novel degradation tags fused to fluorescent proteins. These circuits were tested using both microplate reader and single-cell assays. We first quantified the degradation rates of each tag in isolation. We then tested if there was crosstalk between two distinguishable fluorescent proteins engineered with identical or different degradation tags. We demonstrated that proteolytic crosstalk was indeed not limited to the LAA degradation tag, but was also apparent between other diverse tags, supporting the complexity of the *E. coli* protein degradation system.

## Introduction

Proper cell behavior is maintained using a finite pool of processing resources, such as the limited pool of enzymes required for gene transcription and protein translation^1-2^. Natural biological circuits are largely thought to have evolved to buffer against the effects of limited resources, but we are beginning to understand how processing machinery can form a bottleneck that is in fact leveraged as a control or signaling mechanism^3-4^. Proteolytic (protein degrading) pathways, in particular, have been found to form functional bottlenecks in a native *E. coli* network regulating the stationary phase sigma factor S (σ^S^). The protein σ^S^ is degraded by the ClpXP proteolysis system (ClpXP protease and its chaperones) much faster during exponential growth phase^5-6^ than stationary phase^7^, and the corresponding buildup of σ^S^ in stationary phase acts as a signal triggering the stress response system for starvation^8^. An explanation for increased stability of σ^S^ in stationary phase is that there are an increased number of mistranslated proteins targeted for degradation by ClpXP. Mistranslated proteins are targeted to ClpXP because they have a C-terminal SsrA tag (sometimes labeled LAA tag) due to a special transfer-messenger RNA (tmRNA) being added to mRNA to flag peptides for degradation^9-10^. These proteins compete for a limited number of proteases, especially ClpXP, which results in the formation of a “queue” of substrates for the protease that increases the apparent half-life of σ^S8^.

The complexity of natural proteolytic pathways serves as a barrier to understanding this phenomenon, and so synthetic circuits offer a valuable alternative approach^11^. It was predicted based on the theoretical modeling of a synthetic oscillator that overexpression of LAA-tagged proteins could lead to saturation of proteolytic machinery^12-13^, i.e. that proteolytic machinery could be limiting. Recently, a synthetic circuit more directly demonstrated that two distinguishable fluorescent proteins engineered with ClpXP-targeting LAA-tags can lead the formation of a queue that resulted in crosstalk, such that the buildup of one protein can increase the concentration of another (Fig. 1A)^8^. Queueing theory has since been adopted to describe how competition between substrates for protease can lead to pronounced coupling and statistical correlation^8, 14-16^. The impact of proteolytic queueing competition leads to a rewiring of natural and synthetic circuits to include mutual modulation of substrate degradation rates^17^. This, in particular, applies to all but one existing bacterial synthetic oscillator, which target multiple species of protein to a common protease ClpXP^18^. One exception is the recently modified repressilator^19^, where active degradation by protease was systematically removed to produce a more robust growth-dependent (dilution-dependent) oscillator, which interestingly was predicted based on a prior analysis of proteolytic competition^17^.

**Fig. 1. (A).**
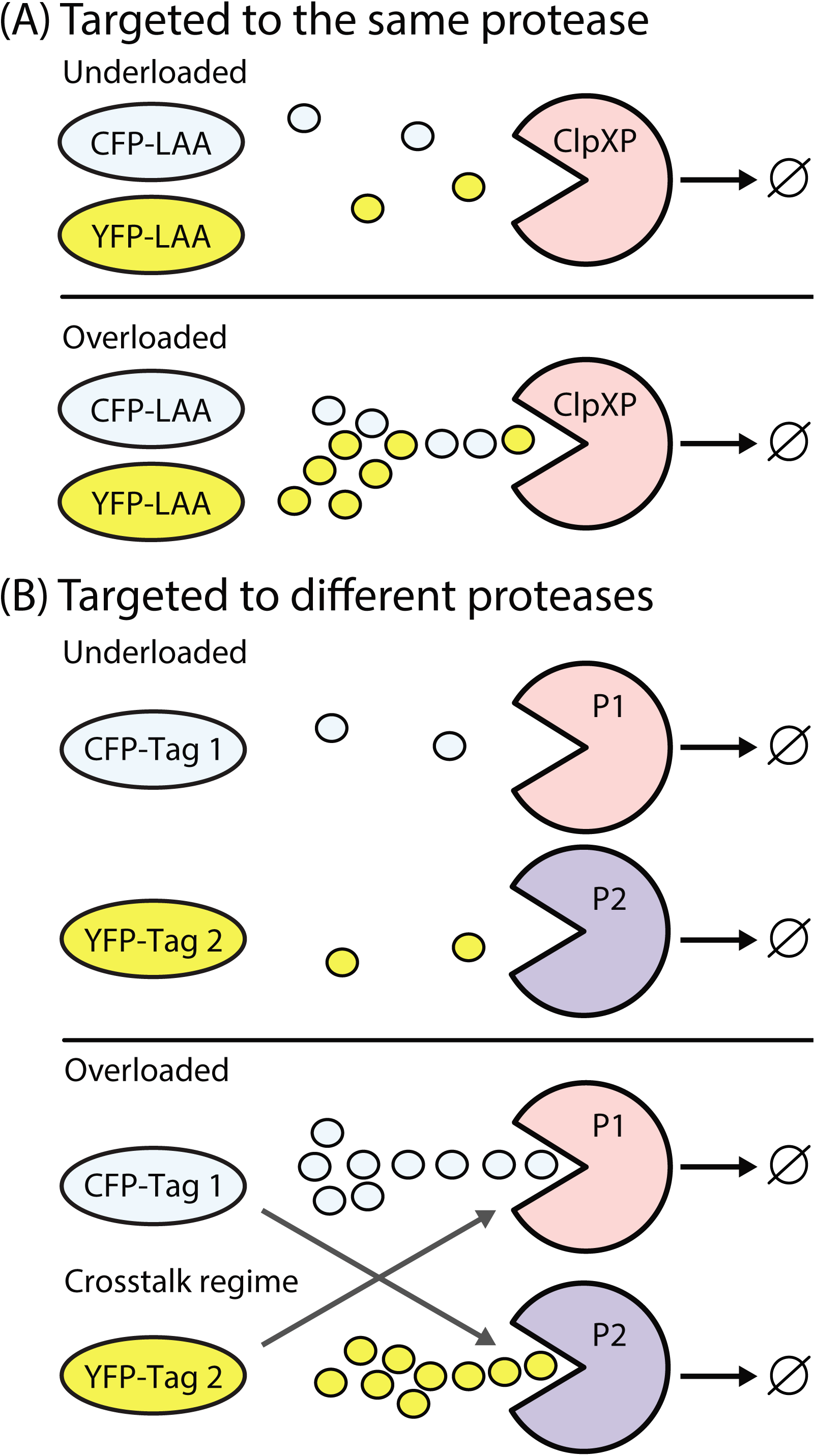
Previous researchers have demonstrated that a queue can form through competition for degradation using fluorescent proteins (CFP and YFP) targeted (through the LAA tag) to the same protease ClpXP^8^. **(B)** We tested if two tags that are specific targets to two different proteases (P1, P2) could form queues.

The single protease crosstalk picture is too simplistic for native circuits, and the reliance on a single degradation pathway for bacterial synthetic oscillators presents a scalability problem that limits the complexity of circuits that can be developed. To address this issue, we investigated the crosstalk between multiple native degradation pathways in *E. coli*. This study extends a prior investigation of computational models that suggested a multi-protease proteolytic bottleneck may still contribute substantially to crosstalk in simple and complex (oscillatory) networks^20^. The influence of crosstalk in multi-protease models was evident even between substrates with substantially different affinities for a protease. Crosstalk may then be generic and complex in native networks. In particular, the different proteolytic networks of *E. coli* may exhibit strong mutual crosstalk, as only three proteases (Lon, ClpXP, and ClpAP, see Table S1) together are required to account for approximately 70%–80% of ATP-dependent degradation in bacterial cells^21-22^. These few proteases thus establish the bulk of proteolytic degradation bandwidth that is shared among a diverse set of actively degraded proteins (about 20% of newly synthesized polypeptides are degraded^23^). This fact combined with the phenomenon of queueing coupling suggests that crosstalk between diverse cellular networks may be typical and cannot be easily relieved by the limited number of proteolytic pathways.

In this work, we used a synthetic biology approach to understand crosstalk between the proteolytic systems of *E. coli*. We designed several synthetic genes that produce fluorescent proteins with protease-targeting tags to serve as probes for and indicators of crosstalk between different protease queues. Using the same *E. coli* strain, we systematically investigated these genes in isolation and in combination. The tags that resulted in substantial degradation in isolation were further investigated for proteolytic crosstalk by co-expressing fluorescent proteins with specific tags. We observed that there is often measurable crosstalk when two fluorescent proteins engineered with identical degradation tags were targeted to the same protease, which is as expected based on the queueing analogy. We also identified a range of crosstalk strengths, ranging from undetectable to high, when proteins are selectively targeted to different protease pathways.

## Results and Discussion

### Multiple synthetic amino acid sequence tags target fluorescent proteins for degradation by different proteases

To study proteolytic degradation by different proteases, fluorescent proteins were engineered with different potential degradation tags on either the N-terminus or C-terminus (Table S2). Previously tested degradation tags could not be compared directly because they were characterized in different *E. coli* strains and under different conditions. To our knowledge, this is the first systematic investigation of multiple degradation tags in *E. coli*. We tested the previously determined degradation tags and several newly designed tags (Table 1A-B). Degradation tags were fused to multiple fluorescent proteins and tested for activity using a high-throughput *in vivo* microplate reader assay. All putative degradation tags tested targeted YFP for degradation (Fig. 2A and Fig. S1). HipBc20, MazE, SoxSn20, RepA15, and HipB were identified as poor degradation tags. In all upcoming experiments, only one poor degradation tag with a medium degradation rate, HipB, was tested for proteolytic crosstalk with other tags. All other tags (LAA, RepA70, MarA, and MarAn20) except SoxS were tested for proteolytic crosstalk.

**Table 1.**
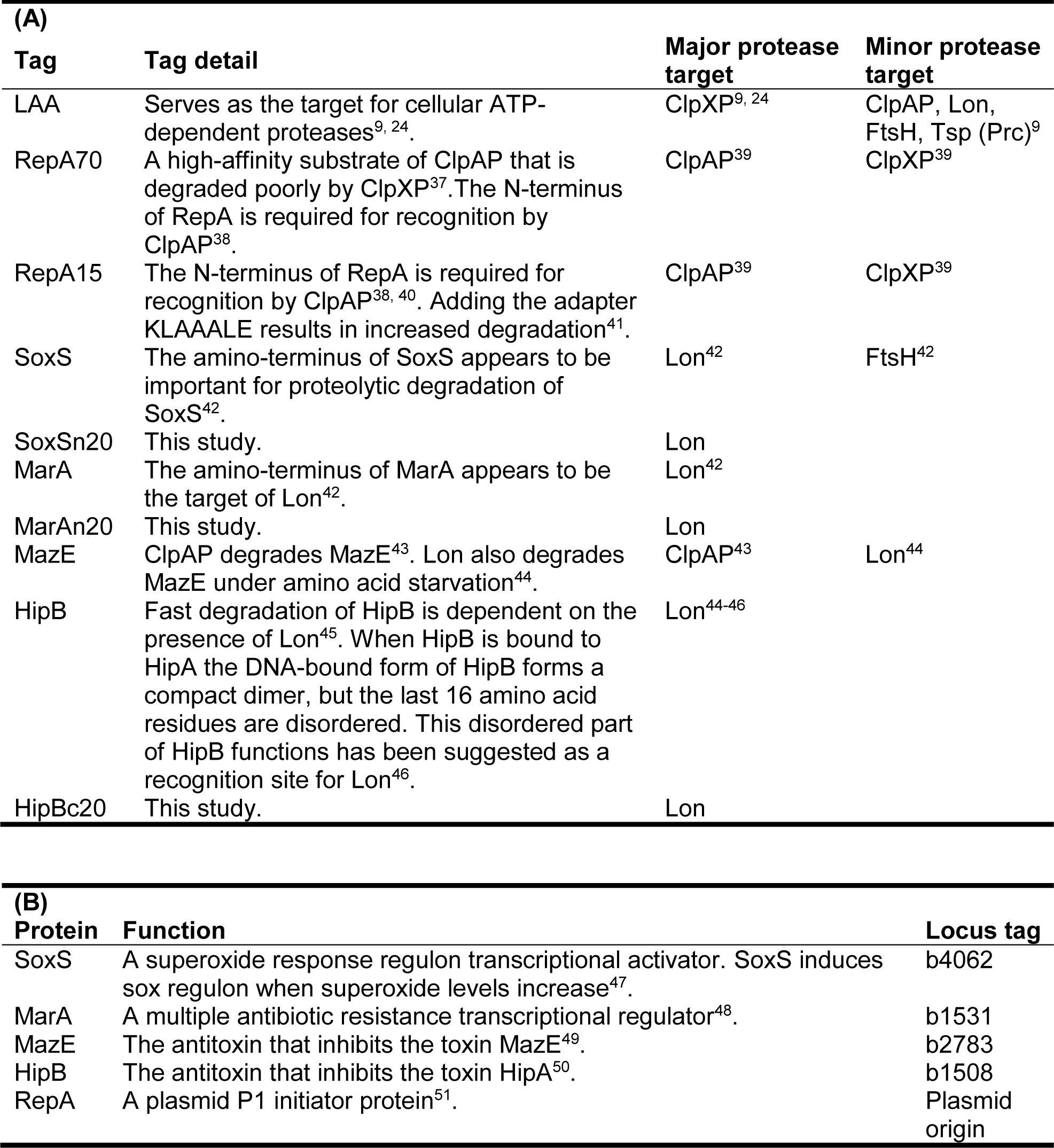
(A) Degradation tags and (B) source (*E. coli* K12 MG1655).

| **(A)** |  |  |  |
| --- | --- | --- | --- |
| **Tag** | **Tag detail** | **Major protease target** | **Minor protease target** |
| LAA | Serves as the target for cellular ATP-dependent proteases<sup>9, 24</sup>. | ClpXP<sup>9, 24</sup> | ClpAP, Lon, FtsH, Tsp (Prc)<sup>9</sup> |
| RepA70 | A high-affinity substrate of ClpAP that is degraded poorly by ClpXP<sup>37</sup>. The N-terminus of RepA is required for recognition by ClpAP<sup>38</sup>. | ClpAP<sup>39</sup> | ClpXP<sup>39</sup> |
| RepA15 | The N-terminus of RepA is required for recognition by ClpAP<sup>38, 40</sup>. Adding the adapter KLAAALE results in increased degradation<sup>41</sup>. | ClpAP<sup>39</sup> | ClpXP<sup>39</sup> |
| SoxS | The amino-terminus of SoxS appears to be important for proteolytic degradation of SoxS<sup>42</sup>. | Lon<sup>42</sup> | FtsH<sup>42</sup> |
| SoxSn20 | This study. | Lon |  |
| MarA | The amino-terminus of MarA appears to be the target of Lon<sup>42</sup>. | Lon<sup>42</sup> |  |
| MarAn20 | This study. | Lon |  |
| MazE | ClpAP degrades MazE<sup>43</sup>. Lon also degrades MazE under amino acid starvation<sup>44</sup>. | ClpAP<sup>43</sup> | Lon<sup>44</sup> |
| HipB | Fast degradation of HipB is dependent on the presence of Lon<sup>45</sup>. When HipB is bound to HipA the DNA-bound form of HipB forms a compact dimer, but the last 16 amino acid residues are disordered. This disordered part of HipB functions has been suggested as a recognition site for Lon<sup>46</sup>. | Lon<sup>44-46</sup> |  |
| HipBc20 | This study. | Lon |  |
| **(B)** |  |  |  |
| **Protein** | **Function** | **Locus tag** |  |
| SoxS | A superoxide response regulon transcriptional activator. SoxS induces sox regulon when superoxide levels increase<sup>47</sup>. | b4062 |  |
| MarA | A multiple antibiotic resistance transcriptional regulator<sup>48</sup>. | b1531 |  |
| MazE | The antitoxin that inhibits the toxin MazE<sup>49</sup>. | b2783 |  |
| HipB | The antitoxin that inhibits the toxin HipA<sup>50</sup>. | b1508 |  |
| RepA | A plasmid P1 initiator protein<sup>51</sup>. | Plasmid origin |  |

**Fig. 2.**
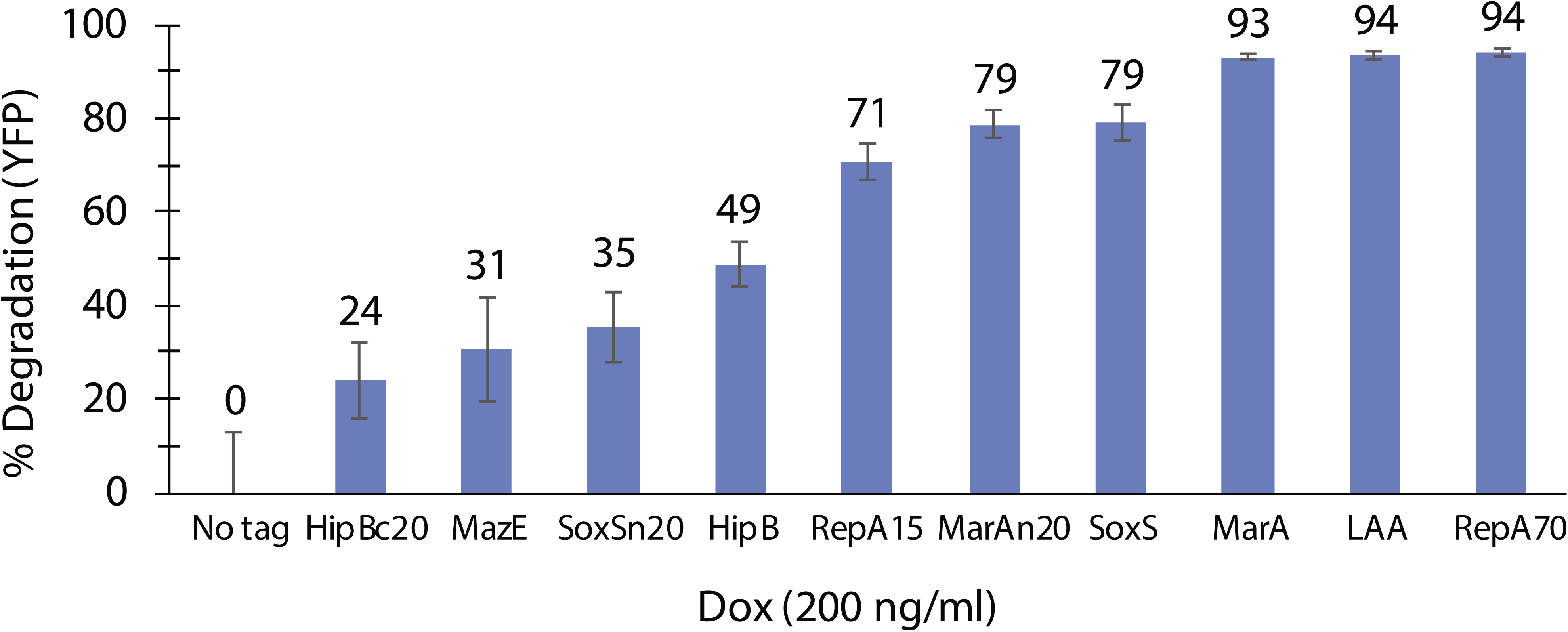
Batch single degradation tag results. YFP derivatives were expressed from the P_LtetO_ promoter using the chemical inducer doxycycline (Dox) at 200 ng/ml. A high-throughput *in vivo* microplate reader assay was used to determine the fluorescent level of proteins with and without degradation tags. The percentage degradation was calculated in this manner: 100% X (1 - YFP_tag_/YFP_untagged_) and standard deviations were calculated using a Taylor expansion^7^. This assumed that untagged YFP is at maximum production and this avoids dividing by a small number. The percentage degradation was calculated for proteins induced at different concentrations of Dox (using data from Fig. S1). Four biological replicas were used to calculate the mean fluorescence and standard deviation from *in vivo* microplate reader batch data.

### The main proteases of an *E. coli* cell can be overloaded when proteins are targeted to a single protease via engineered degradation tags

Previous researchers utilized LAA tagged proteins to demonstrate that proteolytic queues form at ClpXP in the overloaded state ^8^. We recreated their results using our own circuits (Fig. 3B). We then explored if other degradation tags with presumably different affinities (Fig. 1A) would result in the formation of queues and if queues could form with proteases other than ClpXP. YFP and CFP proteins were engineered with RepA70 tags for degradation by ClpAP. The fluorescence levels of RepA70-YFP were monitored as we increase the level of RepA70-CFP proteins by adding the chemical inducer IPTG. The fluorescence of RepA70-YFP increased as more RepA70-CFP was produced (Fig. 3C). This indicated that ClpAP protease can be overloaded, and a proteolytic-queue forms, similar to what was observed with the LAA tagged proteins targeted to ClpXP^8^. We also tested two other tags, MarA and MarA20 (20 amino acids from the N-terminal of MarA), which target proteins to be degraded by the Lon protease. The Lon protease was weakly overloaded by MarA tagged proteins, but was overloaded more by MarAn20 tagged proteins (Fig. 3C-D). This made us wonder if Lon could be overloaded when both MarA and MarAn20 were co-produced. Indeed, this was the case (Fig. 3E).

**Fig. 3.**
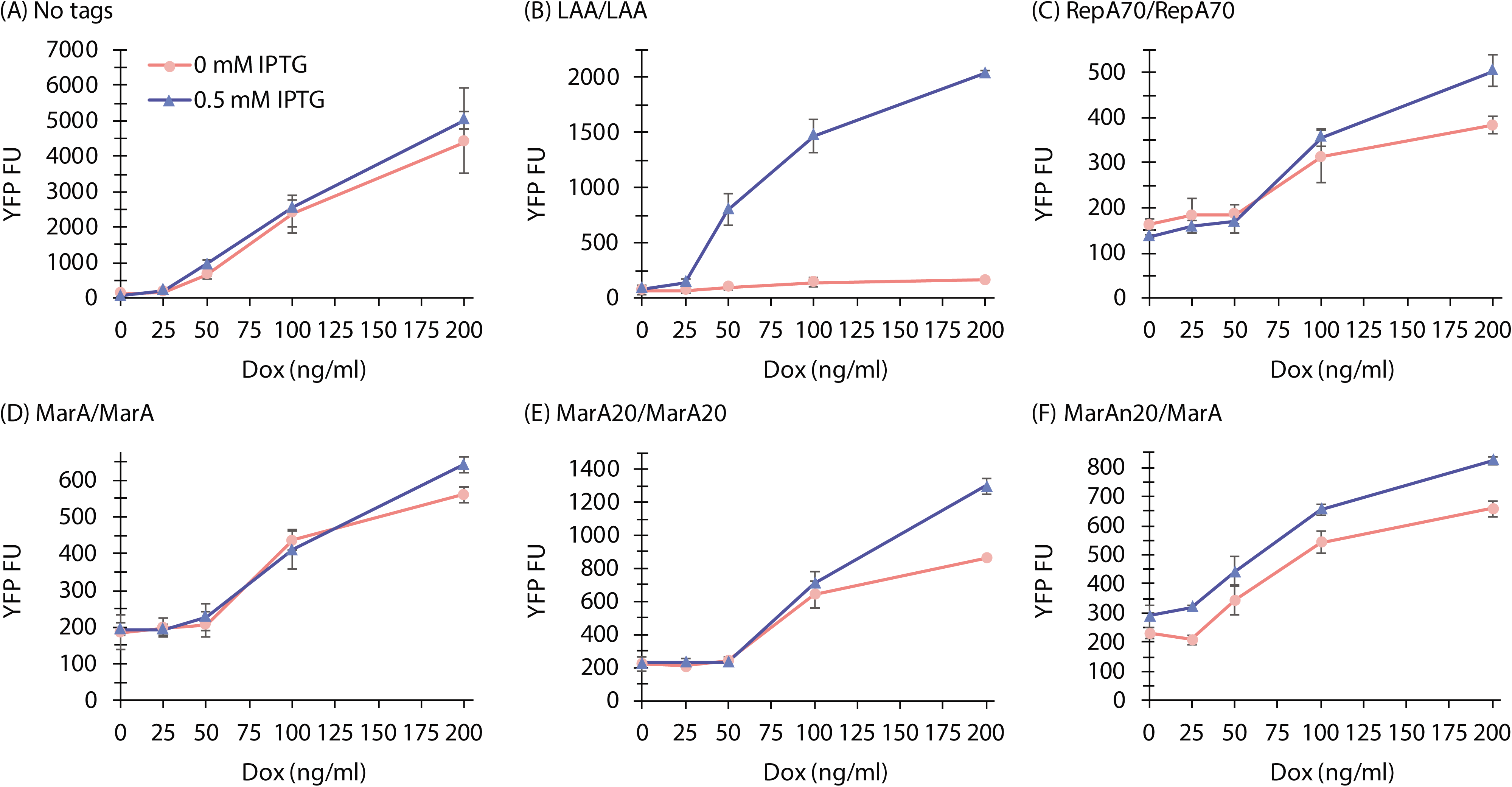
Degradation queues form when the proteins are engineered with degradation tags targeted to the same proteolytic pathway. (**A**) LAA-tagged proteins targeted to ClpXP had the strongest apparent crosstalk, while (**B**) RepA70-tagged proteins target to ClpAP had weak crosstalk compared to what was determined for LAA-tagged proteins. (**C**) MarA tagged proteins targeted to Lon had weak crosstalk, but (**D**) MarAn20-tagged (20 amino acids from the N-terminal of MarA) proteins also targeted to Lon had greater crosstalk. (**E**) Production of CFP-MarAn20 resulted in an increased level of YFP-MarA indicating crosstalk between these two tags targeted to the Lon protease. YFP derivatives were expressed from the P_LtetO_ promoter using the inducer Dox, while CFP derivatives were expressed from P_lac/ara_ promoter using inducer 0.5 mM IPTG (all experiments contained 1% arabinose). Each tag comparison is indicated by tag/tag with CFP being the first tagged and YFP being the second tagged protein. Four biological replicas were used to calculate the mean fluorescence and standard deviation from *in vivo* microplate reader batch data. FU: arbitrary fluorescence unit.

### The main proteases of *E. coli* can exhibit different levels of crosstalk depending on the degradation tags used

We have demonstrated that ClpXP, ClpAP, and Lon can be overloaded using two proteins engineered with identical degradation tags targeted for a specific protease. We have hypothesized that crosstalk between proteases may occur through shared information (Fig. 1B). To test this hypothesis in a synthetic system, we monitored the level of a fluorescent protein (YFP) targeted to one protease while producing another protein (CFP) targeted to a different protease. There was strong crosstalk when proteins with the LAA degradation tag (target to ClpXP) were co-produced with proteins with all other tags (RepA70, MarA, MarAn20, and HipB; Fig. 4A-D). Although ClpXP is the primary protease that degrades LAA-tagged proteins^9, 24^, several other proteases recognize this tag (Table S1). Perhaps due to the importance of removal of potentially harmful waste proteins, cells evolved to have “backup” (a cellular redundancy) proteolytic pathways to remove peptides even if ClpXP is overloaded. Although LAA-tagged proteins are often utilized in synthetic biology circuits^18^, our results suggest it is not an ideal candidate for orthogonal circuits that depend on a proteolytic pathway such as most oscillators. We have tested several other degradation tags that can be potentially used in future synthetic circuits. There was measurable crosstalk when proteins with RepA70 and MarA degradation tags were co-produced (Fig. 4E), but no detectable crosstalk when MarAn20 and RepA70 tagged proteins were co-produced (Fig. 4F). This suggests that MarAn20 and RepA70 may be useful for future synthetic circuits. All proteins with the degradation tags tested thus far were rapidly degraded. We tested a medium-degradation tag, HipB, and, interestingly, we observed no apparent crosstalk between proteins with HipB-tags and all other tagged tested tags (Fig. S2), except for the LAA tag (Fig. 4D).

**Fig. 4.**
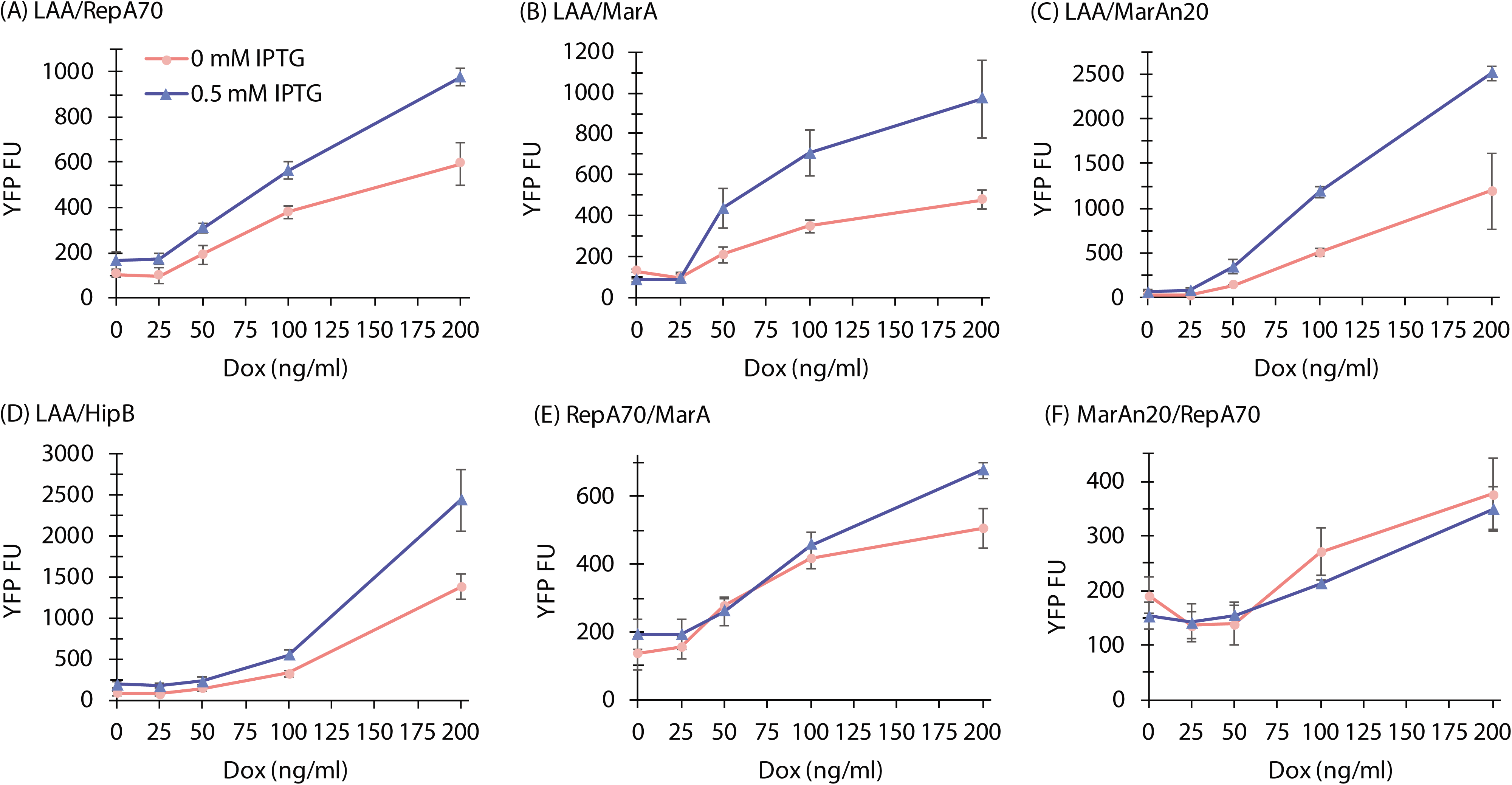
The main proteases of an *E. coli* cell can have different levels of proteolytic crosstalk depending on the degradation tags utilized. Strong crosstalk was detected when proteins with the LAA-tag (targeted to ClpXP) was co-produced with (**A**) RepA70, (**B**) MarA, (**C**) MarAn20, and (**D**) HipB tagged proteins. Proteins engineered with (**E**) RepA70 and MarA tags, and (**F**) MarAn20 and RepA70 tags had measurable crosstalk and no detectable crosstalk, respectively. YFP derivatives were expressed from the P_LtetO_ promoter using the inducer Dox, while CFP derivatives were expressed from P_lac/ara_ promoter using inducer 0.5 mM IPTG (all experiments contained 1% arabinose). Each tag comparison is indicated by tag/tag with CFP being the first tagged and YFP being the second tagged protein. Four biological replicas were used to calculate the mean fluorescence and standard deviation from *in vivo* microplate reader batch data. FU: arbitrary fluorescence unit.

### Single-cell data from microscopy slides support the high-throughput microplate data

The high-throughput microplate batch data allowed us to get an overall picture of the queueing phenomena with a gradient of inducer concentrations, but it is an average measurement of 1000’s of cells. Analysis of single-cell images is an independent technique that is more sensitive than batch data. Single-cell analysis is less subjected to the averaging effects characteristic of batch (population-scale methods) and offers a level of discrete detection that is unobtainable with traditional microbiological techniques^25^. Single-cell data and microplate batch data had similar results when proteins were targeted to the same proteolytic pathways, although crosstalk is more apparent with the single-cell data, especially for the RepA70 tag (Fig. 5A). The single-cell data and microplate batch data also had similar results when proteins were targeted to different pathways (Fig. 5B and Fig. S3). The only minor difference detected by these methods was when proteins with RepA70 and MarA were co-produced. The batch data showed weak crosstalk (crosstalk was only detected at the highest Dox concentration; Fig. 4E), while the single-cell data showed no apparent crosstalk (Fig. 5B).

**Fig. 5.**
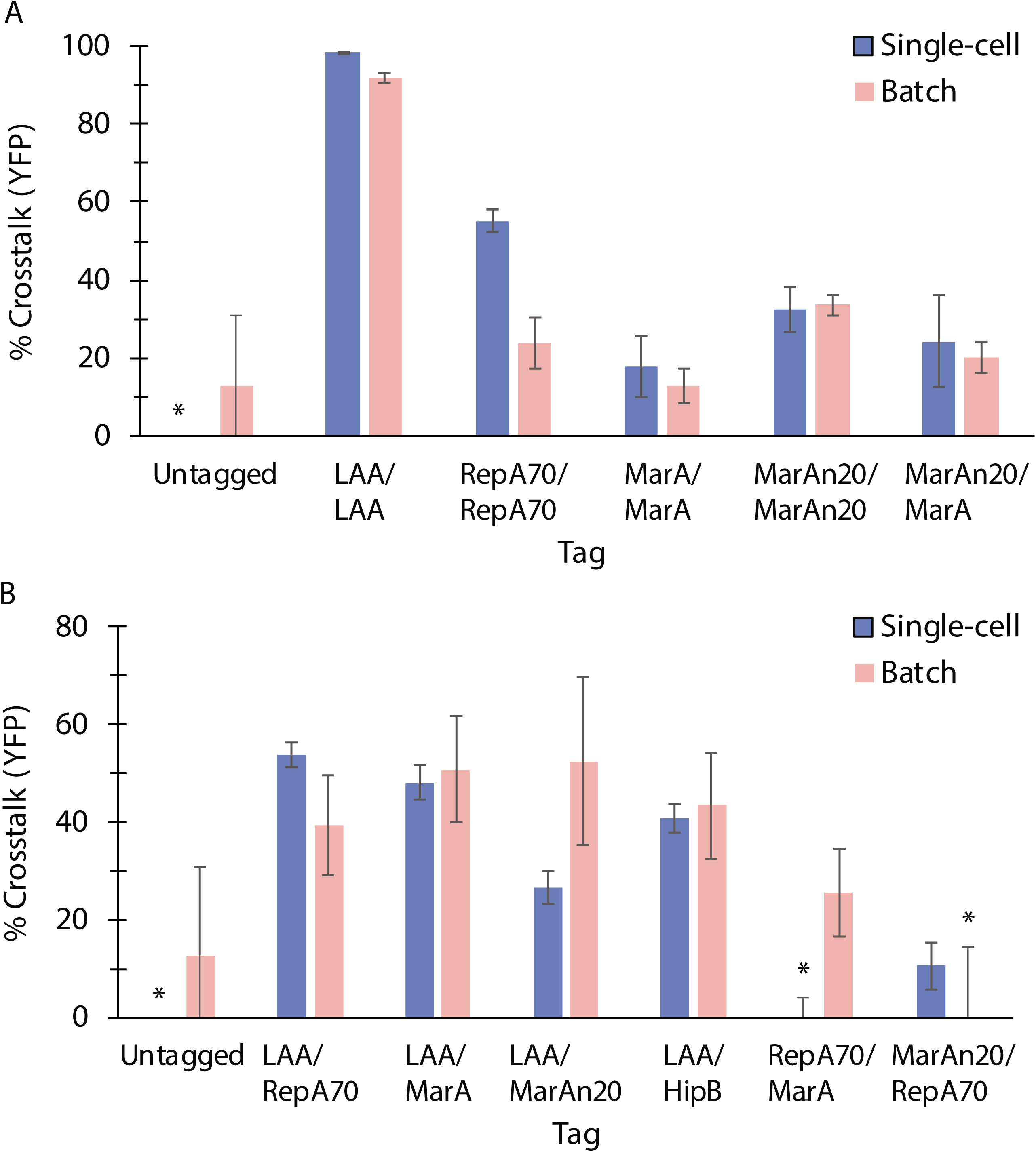
Analysis of crosstalk using both single-cell and batch data. Single-cell data and microplate batch data had similar results with (**A**) proteins engineered with degradation tags targeted to the same protease, and (**B**) proteins engineered with degradation tags targeted to the different proteases. Crosstalk with HipB-tagged proteins was also determined using these methods (Fig. S3). Single-cell data was acquired by imaging cells at 1000X magnification with and without IPTG (inducing CFP derivatives) using a fluorescent confocal microscope. Cells were identified and fluorescence levels were calculated using Fiji, machine learning, and in-house scripts (see Methods). The percentage crosstalk was calculated in this manner: 100%. (1 – YFP_No__IPTG_/YFP_0.5mM__IPTG_) with the standard deviation and SEM calculated using a Taylor expansion ^7^. Use of this ratio avoids dividing by a small number and reduces the impact of statistical error. Each tag comparison is indicated by tag/tag with CFP being the first tagged protein and YFP being the second tagged protein. For each single-cell fluorescent data set, the mean and SEM were calculated from 441-1133 individual cells. For *in vivo* microplate reader batch data, four biological replicas were used to calculate the mean fluorescence and standard deviation. YFP derivatives were expressed from the P_LtetO_ promoter using Dox 200 ng/ml, while CFP derivatives were expressed from P_lac/ara_ promoter using 0.5 mM IPTG (all experiments contained 1% arabinose). *These values are less than (but near) zero.

## Conclusion

Using synthetic circuits, we demonstrated that crosstalk could arise between several different proteolytic pathways. We first characterized a collection of amino acid sequences (tags) that target proteins towards active degradation. Proteins fused to these tags were expressed in a common strain using identical promoters and identical plasmid origins, which allowed us to compare fairly the effectiveness of these tags as degradation signals. This initial characterization of molecular parts is itself of value to synthetic biology. We then co-expressed YFP and CFP with generally different tags to determine crosstalk between pathways. The LAA tag was particularly prone to exhibiting crosstalk with itself and other tags. Other tag combinations demonstrated a range of crosstalk, though not as strong as we measured with LAA. Since many current synthetic systems rely on proteins engineered with LAA tags (targeted to ClpXP)^12, 16, 18, 26-35^, our results strongly suggest that proteolytic crosstalk may be a major hurdle to scalability, indicating that new protease tags are required. Our select pairs of degradation tags with minimal to moderate crosstalk may have future applications in this direction, e.g. in the development of the first synthetic orthogonal to semi-orthogonal oscillators. Of course, tags with strong crosstalk may still be of value, as they could be used for more coordinated modules.

Our findings have implications for both the study of natural systems and the design of complex synthetic systems. In natural systems, cells may use transcriptional regulation and proteolytic coupling as a form of regulation based on the desired response. Transcriptional regulation allows for either a coupled or an uncoupled response. Proteolytic coupling has advantages over a transcriptional response; it may be a quicker mechanism that requires less energy. A transcriptional response to an overloaded protease requires transcription of RNA, production of protein, protein folding, and removal of excess protease after the overabundant substrates are removed. In contrast, proteolytic queueing utilizes other proteases already present in the cell, thus a subsequent response is quick and requires no additional energy to be effective nor to remove the protease later. This means cells can respond to misfolded proteins and the environmental conditions that cause an increase in faulty translation without significantly slowing growth. *E. coli* utilizes proteolytic queues for σ^S^ regulation^8^. Our recent stochastic models suggest that other natural systems such as toxin-antitoxin mechanism used to modulate bacterial persistence may also be augmented by proteolytic queues^20^.

## Methods

### Reagents

All of the reagents were reagent grade and purchased from Sigma-Aldrich, Fisher Scientific, or Thomas Scientific unless otherwise stated.

### Strains and Plasmids

All strains were derived from *E. coli* DH5alphaZ1 (purchased from Dr. Rolf Lutz). All of the plasmid DNA sequences (in GenBank format) are provided in the supporting Information (**Zip file 1**). CFP and YFP gene derivatives were cloned downstream of the P_LtetO_ of p31Cm (chloramphenicol 10 μg/ml)^16^ or downstream of the P_lac/ara_ of p24Km (kanamycin 25 μg/ml). The plasmid p24Km was constructed by PCR amplification and cloning of a T1 terminator into KpnI-ClaI restriction sites of pZE24MCS (purchased from Dr. Rolf Lutz). The CFP and YFP gene derivatives (**Table 1; Table S2**) were purchased from ThermoFisher or GenScript. The cultures were grown in Lysogeny broth (LB).

### Absorbance and fluorescence measurements with a microplate reader

Cells were passed (1:250 dilution) from an overnight culture or a culture stored at 4° C for less than two weeks into LB and antibiotics. Cells were grown at 37° C at a shaking rate of 250 rpm for 2-3 h, and then 100 μl cell culture was added to individual wells of a 96-Well Optical-Bottom Plates with Polymer Base (ThermoFisher) already containing 100 μl of regents (LB, antibiotics, and inducers). A Cytation 3 microplate reader (BioTek) was used to grow and monitor cells. Cell growth was measured at OD_600_ (Optical density at 600 nm). The excitation and emission (Ex/Em) used for YFP and CFP were 510/540 and 447/477 nm, respectively. The wavelengths for Ex/Em were empirically determined to minimize crosstalk between different fluorescent proteins. The background of the media (median absorbance and mean YFP fluorescence) was subtracted from the raw reads. The fluorescence values were compared at OD_600nm_ ∼0.4 (mid-log growth phase) and then the fluorescent values were normalized by dividing by the OD. Four biological replicas were used to calculate the mean fluorescence and standard deviation. Bleed-through from one fluorescent channel into another was tested by two different methods. First, the potential bleed-through from the YFP channel into the CFP channel was tested by producing untagged YFP (the brightest YFP derivative) at different doxycycline (Dox) concentrations (inducing YFP) and monitoring CFP production. A similar test was done with only the untagged CFP protein (induced by isopropyl β-D-1-thiogalactopyranoside (IPTG) and arabinose). No apparent bleed-through was detected in the CFP channel when YFP was produced (Fig. S4A) and no apparent bleed-through was detected in the YFP channel when CFP was produced (Fig. S4B). Second, bleed-through was tested with *E. coli* strains carrying both untagged fluorescent proteins CFP and YFP. No apparent bleed-through was detected in the CFP channel when YFP was produced (Fig. S4C), and no apparent bleed-through was detected in the YFP channel when CFP was produced (Fig. S4D) despite carrying genes for both fluorescent proteins.

### Single-cell snapshots

Cells were harvested from microplate reader wells at mid-log growth phase (near OD_600nm_ 0.4) and single-cell snapshots were taken on a Nikon Ti microscope with CFP and YFP fluorescence cubes at 1000× magnification (a 100× objective coupled to additional 10× magnification). Phase-contrast images and fluorescence images were taken. All fluorescence images were taken with the same exposure (75 ms) and light intensity (6% solo-intensity) based on bleed-through tests. We empirically determined these settings had no apparent bleed-through (Fig. S5).

### Analysis of Mean Fluorescence

To analyze single-cell snapshots, we used a custom pipeline for single-cell segmentation based on machine learning techniques. This process leveraged a FastRandomForest classifier trained and then applied using the Trainable Weka Segmentation tool in Fiji^36^. The classifier used phase contrast images (with pixel values normalized to a common mean and standard deviation) to identify individual cells that were in focus (Fig. S6), and the resulting segmented regions were saved as a separate collection of segmented images. Segmented images were then processed by custom Python 2.7 scripts using the SciPy and OpenCV packages. These scripts both measured single cell fluorescence in corresponding fluorescence images and also computed single cell statistics.

### Model Fitting to Single Tag Microplate Data

Fluorescence data for the single tag constructs in microplate experiments were fit to a simple enzymatic degradation model in order to describe the data points by a mathematical function. We checked whether the measured YFP-tag fluorescence *Y*_*T*_ (for a range of tags *T*) for a given concentration [*DOX*] could be described by the steady-state of the ODE:

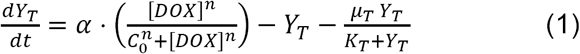

i.e., with *Y*_*T*_ the solution to the equation

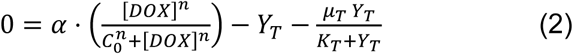

In this model, we have rescaled time such that the dilution rate is 1.0, and we allow the apparent enzymatic velocity *µ*_*T*_ and molar constant *K*_*T*_ to, in principle, have independent values for each tag. The parameters α and *C*_0_ characterize the production rate of protein for a given level of [*DOX*]. We assume that untagged YFP proteins are degraded with zero enzymatic velocity, i.e. *μ*_*Untagged*_ = 0. All data points, excepting data for YFP-RepA15, were jointly fit to this model using the scipy.optimize.minimize function from the SciPy library. Parameter values for one of our best fits are reported in SI Table S3, and the fit is displayed graphically in SI Fig. S1. It is worth noting that if fitted value for *K*_*T*_ is large relative to typical values of *Y*_*T*_, then the relevant degradation parameter is the first-order degradation rate constant, *μ*_*T*_ /*K*_*T*_, and only this ratio is likely very meaningful regarding model fit.

## Acknowledgements

Funding for this research was provided by the 454 National Science Foundation under Grant No. MCB-1330180.

## Supporting Information Tables and Captions

**Table S1.**
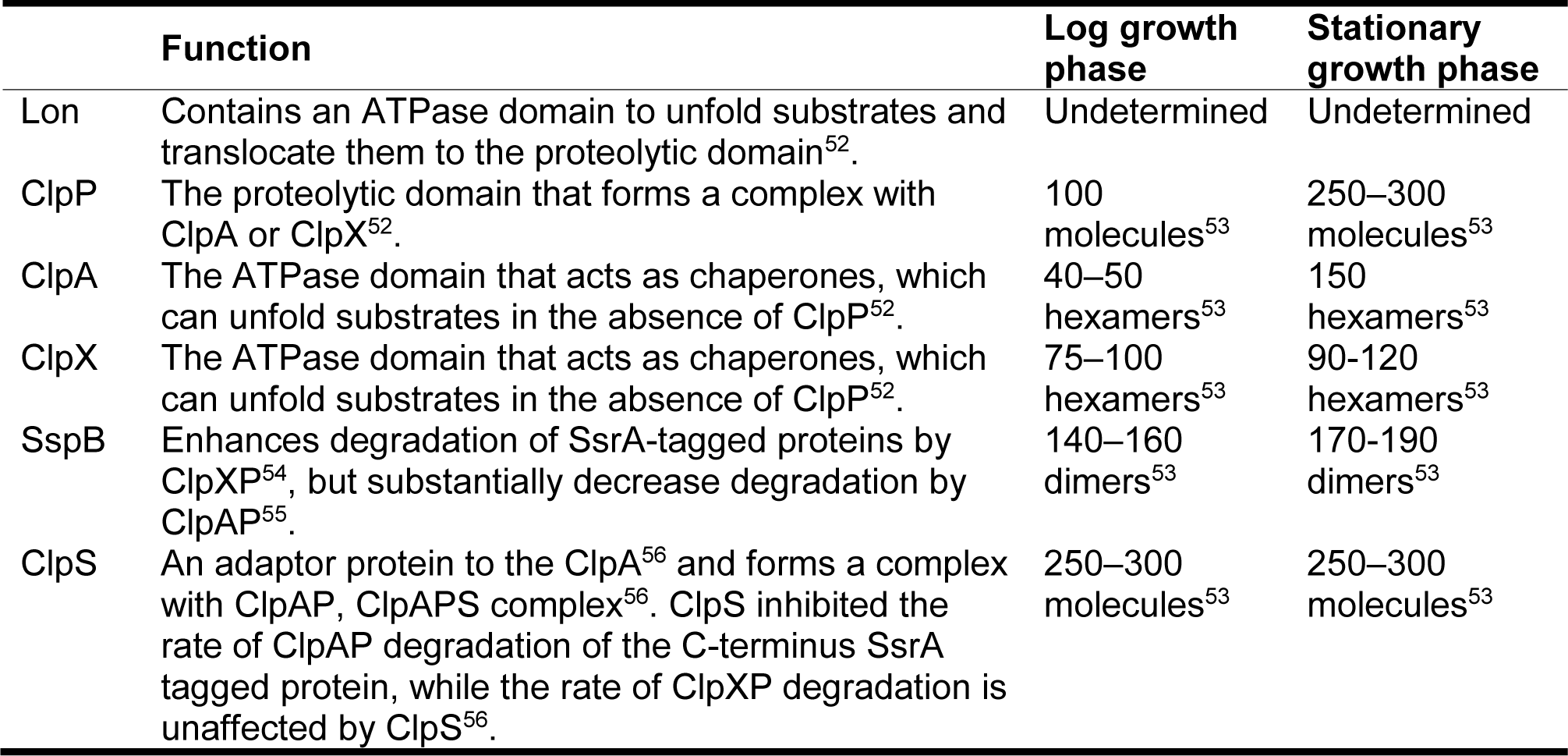
Key cytoplasmic proteases and chaperones.

**Table S2.** Plasmids and strains. (**A**) The parent plasmids were either p31Cm or p24Km plasmid. (**B**) *E. coli* strains were derived from DH5alphaZ1. Arabinose + IPTG, and Dox induce genes under the control of the P_lac/ara_ and P_LtetO_ promoters, respectively. aa: amino acids.

| (A) |  |  |  |  |
| --- | --- | --- | --- | --- |
| Plasmid | Promoter | Tag | Tag terminal | Tag detail |
| p31Cm | $P_{LtetO}$ | None | Untagged | |
| p31CmNB02 | $P_{LtetO}$ | YFP | Untagged | |
| p31CmNB95 | $P_{LtetO}$ | YFP-LAA | C | 11 aa LAA tag |
| p31CmNB44 | $P_{LtetO}$ | YFP-RepA15 | C | 15 aa from the N-terminal of RepA15 with KLAAALE linker |
| p31CmNB04 | $P_{LtetO}$ | RepA70-YFP | N | 70 aa from the N-terminal of RepA |
| p31CmNB88 | $P_{LtetO}$ | YFP-SoxS | C | SoxS with TS linker |
| p31CmNB89 | $P_{LtetO}$ | YFP-SoxSn20 | C | 20 aa from the N-terminal of SoxS with TS linker |
| p31CmNB90 | $P_{LtetO}$ | YFP-MarA | C | MarA with TS linker |
| p31CmNB91 | $P_{LtetO}$ | YFP-MarAn20 | C | 20 aa from the N-terminal of MarA with TS linker |
| p31CmNB92 | $P_{LtetO}$ | YFP-MazE | C | MazE with TS linker |
| p31CmNB93 | $P_{LtetO}$ | YFP-HipB | C | HipB with TS linker |
| p31CmNB94 | $P_{LtetO}$ | YFP-HipBc20 | C | 20 aa from the C-terminal of HipB with TS linker |
| p24Km | $P_{lac/ara}$ | None | Untagged | |
| p24KmNB83 | $P_{lac/ara}$ | CFP | Untagged | |
| p24KmNB82 | $P_{lac/ara}$ | CFP-LAA | C | 11 aa LAA tag |
| p24KmNB07 | $P_{lac/ara}$ | RepA70-CFP | N | 70 aa from the N-terminal of RepA |
| p24KmNB75 | $P_{lac/ara}$ | CFP-SoxS | C | SoxS with TS linker |
| p24KmNB76 | $P_{lac/ara}$ | CFP-SoxSn20 | C | 20 aa from the N-terminal of SoxS with TS linker |
| p24KmNB77 | $P_{lac/ara}$ | CFP-MarA | C | MarA with TS linker |
| p24KmNB78 | $P_{lac/ara}$ | CFP-MarAn20 | C | 20 aa from the N-terminal of MarA with TS linker |
| p24KmNB79 | $P_{lac/ara}$ | CFP-MazE | C | MazE with TS linker |
| p24KmNB80 | $P_{lac/ara}$ | CFP-HipB | C | HipB with TS linker |
| p24KmNB81 | $P_{lac/ara}$ | CFP-HipBc20 | C | 20 aa from the C-terminal of HipB with TS linker |

| **(B)** |  |  |  |
| --- | --- | --- | --- |
| **Strain** | **Plasmids** | **P<sub>lac/ara</sub> promoter** | **P<sub>LtetO</sub> promoter** |
| DZ66 | p24KmNB82, p31CmNB95 | CFP-LAA | YFP-LAA |
| DZ67 | p24KmNB07, p31CmNB95 | RepA70-CFP | YFP-LAA |
| DZ68 | p24KmNB77, p31CmNB95 | CFP-MarA | YFP-LAA |
| DZ69 | p24KmNB78, p31CmNB95 | CFP-MarAn20 | YFP-LAA |
| DZ70 | p24KmNB80, p31CmNB95 | CFP-HipB | YFP-LAA |
| DZ72 | p24KmNB82, p31CmNB04 | CFP-LAA | RepA70-YFP |
| DZ33 | p24KmNB07, p31CmNB04 | RepA70-CFP | RepA70-YFP |
| DZ73 | p24KmNB77, p31CmNB04 | CFP-MarA | RepA70-YFP |
| DZ74 | p24KmNB78, p31CmNB04 | CFP-MarAn20 | RepA70-YFP |
| DZ75 | p24KmNB80, p31CmNB04 | CFP-hipB | RepA70-YFP |
| DZ77 | p24KmNB82, p31CmNB90 | CFP-LAA | YFP-MarA |
| DZ78 | p24KmNB07, p31CmNB90 | RepA70-CFP | YFP-MarA |
| DZ79 | p24KmNB77, p31CmNB90 | CFP-MarA | YFP-MarA |
| DZ80 | p24KmNB78, p31CmNB90 | CFP-MarAn20 | YFP-MarA |
| DZ81 | p24KmNB80, p31CmNB90 | CFP-HipB | YFP-MarA |
| DZ83 | p24KmNB82, p31CmNB91 | CFP-LAA | YFP-MarAn20 |
| DZ84 | p24KmNB07, p31CmNB91 | RepA70-CFP | YFP-MarAn20 |
| DZ85 | p24KmNB77, p31CmNB91 | CFP-MarA | YFP-MarAn20 |
| DZ86 | p24KmNB78, p31CmNB91 | CFP-MarAn20 | YFP-MarAn20 |
| DZ87 | p24KmNB80, p31CmNB91 | CFP-HipB | YFP-MarAn20 |
| DZ89 | p24KmNB82, p31CmNB93 | CFP-LAA | YFP-hipB |
| DZ90 | p24KmNB07, p31CmNB93 | RepA70-CFP | YFP-hipB |
| DZ91 | p24KmNB77, p31CmNB93 | CFP-MarA | YFP-hipB |
| DZ92 | p24KmNB78, p31CmNB93 | CFP-MarAn20 | YFP-hipB |
| DZ93 | p24KmNB80, p31CmNB93 | CFP-HipB | YFP-hipB |

**Table S3.**
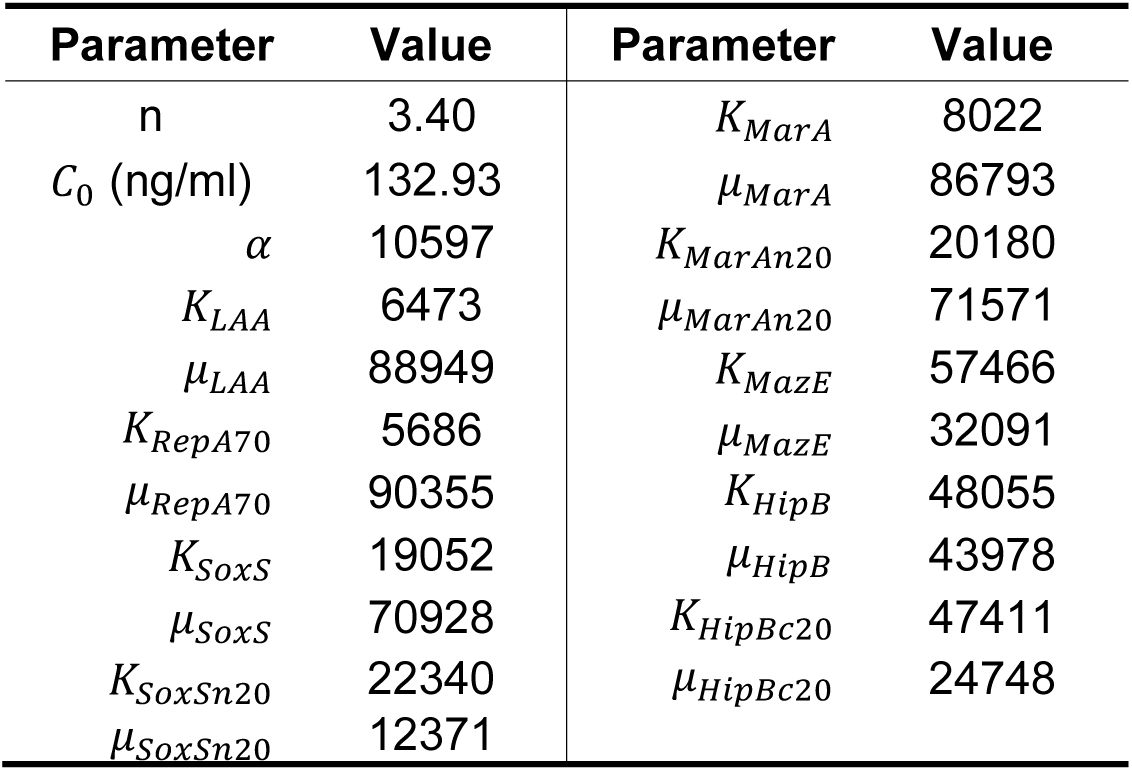
Extracted parameters for single tag fluorescence data. A simple model based on enzymatic degradation was fit to fluorescence data to produce smooth curves that fit through the data. The parameters producing the best fit for this model are contained in this table. These parameters are used in SI Fig S1. Interestingly, the *K* values for the most rapidly degrading species (LAA, RepA70, and MarA) are the smallest estimated values for *K*, suggesting higher affinity enzymatic degradation for these species. All parameters are dimensionless or arbitrary units, excepting for the constant *C*_0_.

| Parameter | Value | Parameter | Value |
| --- | --- | --- | --- |
| n | 3.40 | $K_{MarA}$ | 8022 |
| $C_0$ (ng/ml) | 132.93 | $\mu_{MarA}$ | 86793 |
| $\alpha$ | 10597 | $K_{MarAn20}$ | 20180 |
| $K_{LAA}$ | 6473 | $\mu_{MarAn20}$ | 71571 |
| $\mu_{LAA}$ | 88949 | $K_{MazE}$ | 57466 |
| $K_{RepA70}$ | 5686 | $\mu_{MazE}$ | 32091 |
| $\mu_{RepA70}$ | 90355 | $K_{HipB}$ | 48055 |
| $K_{SoxS}$ | 19052 | $\mu_{HipB}$ | 43978 |
| $\mu_{SoxS}$ | 70928 | $K_{HipBc20}$ | 47411 |
| $K_{SoxSn20}$ | 22340 | $\mu_{HipBc20}$ | 24748 |
| $\mu_{SoxSn20}$ | 12371 | | |

## Supporting Information Captions

**Fig. S1. Single degradation tag results with different levels of Dox.** An *in vivo* microplate reader assay was used to determine the fluorescence of YFP proteins with and without degradation tags. Symbols (connected by dashed lines) represent the mean fluorescence from four biological replicates, and error bars are the calculated standard deviations from these replicates. Solid lines represent fits to a simple mathematical model (see Methods), with parameters given in Table S3. SoxSn20 tagged proteins deviated noticeably from our model fit, but were included in our fitting analysis. RepA15 tagged proteins were excluded from our model fit due to weak degradation and irregular response, though we still report the corresponding percent degradation in Fig. 2.

**Fig. S2. Proteins engineered with the HipB tag and co-produced with (C) RepA70, (D) MarA, and (E) MarAn20 tagged proteins had no detectable crosstalk.** YFP derivatives were expressed from the P_LtetO_ promoter using the inducer Dox; while CFP derivatives were expressed from P_lac/ara_ promoter using 0.5 mM IPTG (all experiments contained 1% arabinose). Each tag comparison is indicated by tag/tag with CFP being the first tagged protein and YFP being the second tagged protein. Four biological replicas were used to calculate the mean fluorescence and standard deviation from *in vivo* microplate reader batch data. FU: arbitrary fluorescence unit.

**Fig. S3. Analysis of crosstalk using both single-cell and batch data with HipB-tagged proteins.** See Fig. 5 and the Method section for details on acquisition and analysis of the single-cell data and microplate batch data.

**Fig. S4. Test for bleed-through from one channel to another in *in vivo* microplate reader experiments.** No apparent bleed-through was detected in the (**A**) CFP channel when YFP was produced and no apparent bleed-through was detected in the (**B**) YFP channel when CFP was produced. Bleed-through was tested with the *E. coli* strains carrying both untagged fluorescent proteins CFP and YFP. (**C**) No apparent bleed-through was detected in the CFP channel when YFP was produced. (**D**) No apparent bleed-through was detected in the YFP channel when CFP was produced. Dox was used to induce YFP expression and IPTG (with 1% arabinose) was used to induce CFP expression. Four biological replicas were used to calculate the mean fluorescence and standard deviation. FU: arbitrary fluorescence unit.

**Fig. S5. Test for bleed-through from one channel to another in single-cell snapshot experiments.** No apparent bleed-through was detected in the CFP channel when YFP was produced and no apparent bleed-through was detected in the YFP channel when CFP was produced. Bleed-through was tested with the *E. coli* strains carrying no fluorescent proteins and untagged fluorescent proteins CFP and YFP. When YFP was produced the CFP channel had no apparent change indicated; the CFP channel when YFP had similar mean fluorescence as the CFP channel with no fluorescent proteins (the mean values were within 1 SEM). The same was true when the CFP was produced and the YFP channel was monitored. Dox 200 ng/ml was used to induce YFP expression and 0.5 mM IPTG + 1% arabinose was used to induce CFP expression. Four biological replicas were used to calculate the mean fluorescence and standard deviation. FU: arbitrary fluorescence unit. Single-cell data was acquired by imaging cells at 1000X magnification using a fluorescent confocal microscope with an exposure of 75 ms and light intensity of 6% for both the CFP and YFP channel. Cells were identified and fluorescence levels were calculated using Fiji, machine learning, and in-house scripts (see Methods). The mean and SEM were calculated from 10-19 individual cells.

**Fig. S6. Method for Single Cell Analysis.** (**a**) To extract single cell fluorescence data, phase contrast images (shown) were taken in sequence with fluorescence images. Phase contrast images generally contained both cells in focus and out of focus. (**b**) Machine learning-based classification (see Methods) was applied to phase contrast images to identify individual cells that were in focus (cells indicated by colored regions). The regions in the image associated with individual cells were then used to measure fluorescence in corresponding images.

